# ATAC-seq of Low-Input and Cryopreserved Primordial Germ Cells Reveals Functional Enhancers

**DOI:** 10.1101/2025.08.30.672031

**Authors:** Akane Kawaguchi, Mao Igari, Yasuto Murayama, Hiroko Iikawa, Mika Sakamoto, Yasukazu Nakamura, Shigehiro Kuraku, Keisuke Ishihara, Daisuke Saito

## Abstract

Cell fate specification during development is orchestrated by dynamic changes in gene expression, driven by the accessibility of cis-regulatory elements. To profile such functional regulatory elements at the genome-wide level, bulk ATAC-seq is a powerful method. Yet, its application has still been technically limited in rare cell types. Here, we optimized and titrated the ATAC-seq protocol for cultured chicken primordial germ cells (PGCs) and obtained high-quality chromatin accessibility profiles from as few as 200 cells, even after cryopreservation. Integrative analysis of our ATAC-seq with published RNA-seq and tissue ATAC-seq datasets identified over 10,000 PGC-specific accessible chromatin regions (ACRs), many absent from somatic tissues. Reporter assays demonstrated activity of selected enhancers in cultured PGCs, while transcriptome integration revealed preferential association of PGC-specific ACRs with genes highly expressed at embryonic day 2.5. *In vivo* transplantation confirmed that these enhancers are active in an early-stage-specific manner, but become silenced upon gonadal settlement. In conclusion, this streamlined method provides a practical and cost-effective approach to the epigenomic analysis of scarce cell populations, facilitating future investigations in developmental biology, germline research, and beyond.

## Introduction

Cell fate specification and differentiation during development are governed by tightly regulated, dynamic changes in gene expression. Increasing evidence indicates that such transcriptional shifts are orchestrated through changes in chromatin accessibility at cis-regulatory elements. Therefore, capturing the chromatin landscape in a spatiotemporal manner is essential to understand and manipulate lineage decisions.

ATAC-seq (Assay for Transposase-Accessible Chromatin using sequencing) enables genome-wide profiling of accessible chromatin with high resolution using far fewer input materials than other conventional profiling methods, such as the DNase I hypersensitivity assay (Ling and Waxman, 2013), allowing profiling from as few as 500 freshly isolated cells (Buenrostro *et al*., 2013). The Omni-ATAC-seq protocol has further improved signal-to-noise ratios through simplified procedures and the use of additional solvents, and has successfully reduced mitochondrial DNA (mtDNA) contamination (Corces *et al*., 2017). However, applying this method to rare embryonic cells remains challenging, as viable cell isolation is technically demanding and often yields limited material. Cells or tissue cryopreservation could address these issues (Fujiwara *et al*., 2019; Halstead *et al*., 2020; Peng *et al*., 2021), but robust ATAC-seq protocols for frozen low-input samples are still underdeveloped. Although single-cell ATAC-seq offers an alternative (Buenrostro *et al*., 2015), its cost and complexity limit its general accessibility (Pott and Lieb, 2015).

Primordial germ cells (PGCs) are an inherently small cell population in developing embryos. They originate outside the embryo proper and migrate dynamically to the gonads, where they undergo sex-specific programs of mitosis, meiosis, stem cell maintenance, and eventual differentiation into gametes (Wear, McPike and Watanabe, 2016; Yoshida, 2020; Barton *et al*., 2024). Thus, changes in chromatin state within the germline are of broad biological interest, and substantial knowledge has been accumulated regarding DNA and histone modifications (Seki *et al*., 2007; Nagano *et al*., 2022). However, compared to these epigenetic marks, much less is known about the open–closed chromatin status of PGCs as revealed by ATAC-seq. Furthermore, because PGCs are known to have relatively high mitochondrial content (Motta *et al*., 2000; Deniz Koç and Yüce, 2012), they provide an excellent benchmark model for evaluating and optimizing ATAC-seq protocols with respect to mitochondrial genome contamination—a common technical challenge in this assay.

In this study, we leveraged the unique advantages of cultured chicken PGCs and the chicken embryo model. Cultured PGCs can be expanded *in vitro* while preserving germline potential and an *in vivo*–competent migratory program, homing to the gonads via the endogenous route after transplantation (van de Lavoir *et al*., 2006; Whyte *et al*., 2015; Morimoto and Saito, 2025). These properties enable rigorous benchmarking of ATAC-seq protocols and functional testing of cis-regulatory elements within their physiological context.

Here, we first optimized the Omni-ATAC-seq workflow for low-input and cryopreserved chicken PGCs, demonstrating that high-confidence chromatin accessibility profiles can be obtained from as few as 200 cells, even after freezing. This optimization also revealed sequence biases of Tn5 insertions in the chicken genome, underscoring the need for proper background controls. We then established an *in vitro* PGC differentiation system within this study, enabling a reporter assay to functionally evaluate PGC-specific and stage-specific enhancers identified from ATAC-seq data. Using transplantation into chicken embryos, we further demonstrated that selected enhancers are active in migrating PGCs *in vivo*. Together, these results show that our approach enables robust identification of functional cis-regulatory elements from minimal starting material, providing a powerful framework for studying chromatin regulation not only in germline biology but also in other developmentally dynamic or clinically relevant rare cell types.

## Results and discussions

### Cryopreservation impacts mtDNA content but not overall library quality

To evaluate the impact of cell number and cryopreservation on Omni-ATAC-seq quality, we prepared cultured chicken PGC samples with four different input sizes (200, 1,000, 5,000, and 25,000 cells), using either freshly harvested or cryopreserved cells stored at –80°C for one week (Fig. 1A). The protocol followed the original Omni-ATAC-seq method (Corces *et al*., 2017), which employs a combination of detergents (Tween-20, NP-40, Digitonin) for effective membrane permeabilization, and PBS addition during tagmentation to enhance the signal-to-noise ratio by reducing mtDNA contamination. Cryopreserved samples exhibited high cell viability (>90%) based on trypan blue exclusion, and nuclear morphology remained comparable to that of live cells, as assessed by nuclear staining (Fig. 1B, C). These results indicate that both fresh and cryopreserved cells were suitable as starting materials. As a negative control, we included a sample of purified genomic DNA subjected to tagmentation. After library construction and sequencing, we processed fastq files through standard pre-processing steps, including adapter trimming, filtering, genome alignment, and duplicate removal. Cryopreserved samples exhibited higher duplication rates (19–27%) compared to live cells (9–14%) (Fig. 1D, Supplementary Table 1), suggesting that freeze–thaw steps may increase redundant amplification.

**Figure 1.**
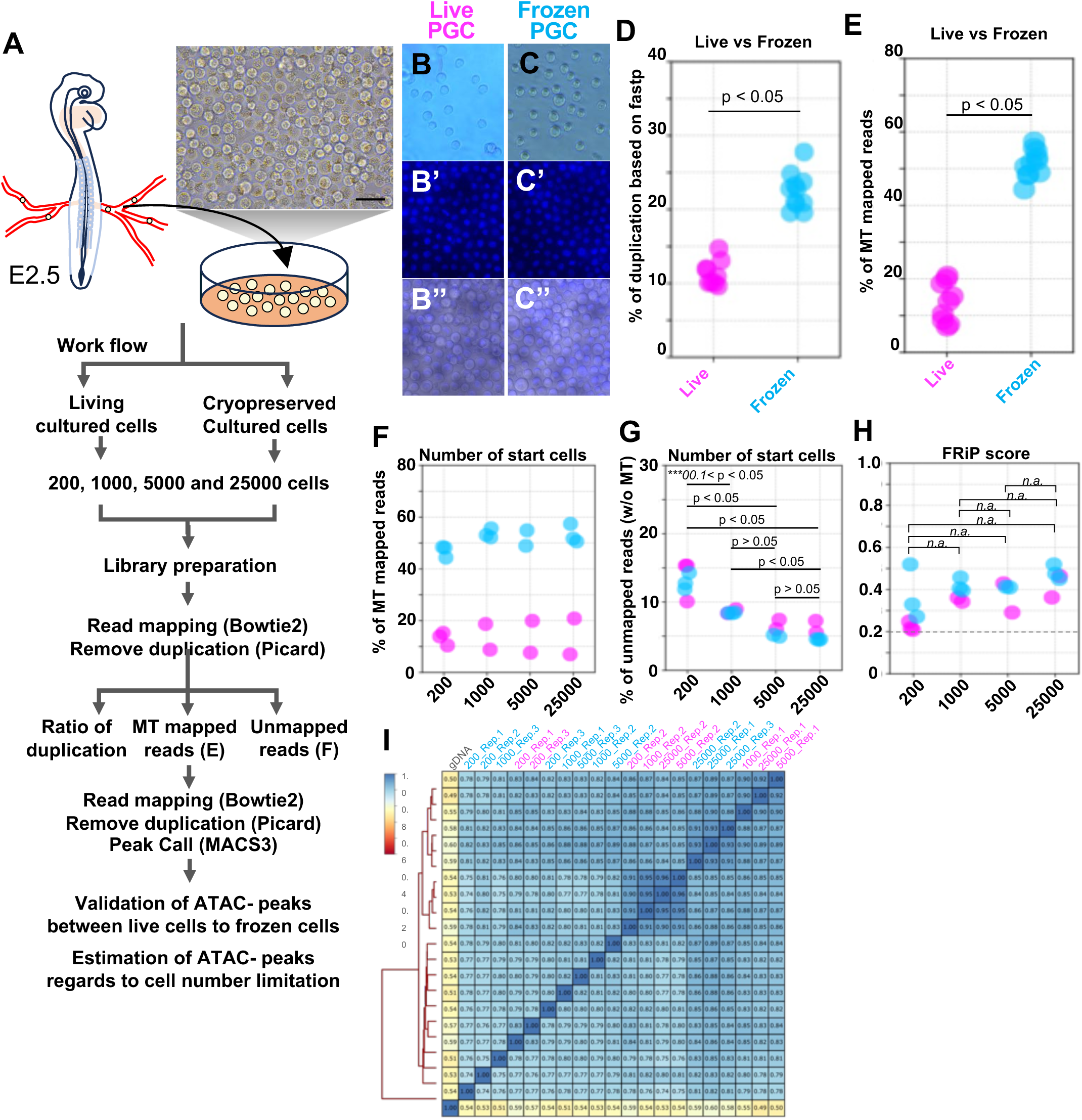
Quality assessments for Omni-ATAC-seq of live and frozen cultured PGCs. (A) Schema showed the harvesting of PGCs from E2.5 chicken embryos and the workflow of the Omni-ATAC-seq benchmarking setup. (B–C) Cell viability analysis using NucBlue staining in live PGCs (B–B″) and frozen PGCs (C–C″). Frozen PGCs retained nuclear morphology comparable to that of live cells. (D–H) Quality metrics across input conditions, (D) duplication rates detected after initial quality control and adapter trimming using Fastp, and (E) proportion of unmapped reads after alignment. (F-G) contamination rates of mitochondrial reads. (H) FRiP score for the individual libraries. In E to I, live and frozen samples marked with magenta and blue, respectively. (I) Heatmap comparing all Omni-ATAC-seq libraries across a range of input cell numbers for both live and frozen conditions.

We next evaluated three key quality metrics of ATAC-seq data: (i) mtDNA (mtDNA) mapping rate, (ii) unmapped read rate, and (iii) FRiP (Fraction of Reads in Peaks) score. The mtDNA read contamination was notably higher in cryopreserved samples (50–60%) than in fresh samples (5–15%) (Fig. 1E, F), suggesting that cryopreservation may reduce the efficiency of mitochondrial removal during lysis. This observation is consistent with earlier reports on ATAC-seq protocols lacking detergent-based optimization (Fujiwara *et al*., 2019). The proportion of unmapped reads was highest in 200-cell input samples (10–15%), regardless of cryopreservation, while samples with ≥1,000 cells showed stable mapping rates (∼5–7%) (Fig. 1G). This suggests that input cell number, rather than freeze–thaw condition, is the major determinant of read mappability. FRiP scores, a standard measure of ATAC-seq signal quality, exceeded 0.2 in all libraries (Fig. 1H), thereby meeting ENCODE guidelines (https://www.encodeproject.org/atac-seq/). While FRiP scores showed a weak correlation with input cell number, they were not significantly influenced by cryopreservation. In the end, after removing duplicate reads and mtDNA-mapped reads, the consistent correlation signatures obtained from the BAM files do not exhibit large differences across conditions (Fig. 1H). Taken together, these results demonstrate that while cryopreservation increases mtDNA contamination and duplication rates, it does not substantially compromise library quality— even in PGC samples, which are known to contain relatively high levels of mtDNA (Motta *et al*., 2000; Deniz Koç and Yüce, 2012). Importantly, reproducible and high-quality chromatin accessibility profiles can be obtained from as few as 200 cells, even after cryopreservation.

### Peak Number and Sensitivity in Low-Input Cryopreserved Samples

We next examined how variations in input cell number and cryopreservation influence the number and quality of peaks detected by ATAC-seq. To begin, peak calling was performed using MACS3 without normalization (–broad, q < 0.05). In live-cell libraries, the number of identified peaks ranged from 31,018 to 48,786. In contrast, cryopreserved libraries yielded significantly fewer peaks, between 22,737 and 38,558, with a statistically significant reduction compared to live samples (*P* < 0.05; Fig. 2A, Supplementary Table 1). Peak counts showed a modest upward trend with increasing input cell numbers in both conditions (*P* < 0.05; Fig. 2B). To assess whether these differences stemmed from sequencing depth, we next normalized all libraries to 10 million valid reads using computed scaling factors (Supplementary Table 2). After reading subsampling and re-calling peaks with identical MACS3 parameters, peak numbers across all libraries converged, with no statistically significant differences between groups (P > 0.05), except for the gDNA-derived negative control (Fig. 2C). It suggests that the reduced peak counts in cryopreserved samples primarily reflect lower sequencing depth rather than intrinsic biological differences.

**Figure 2.**
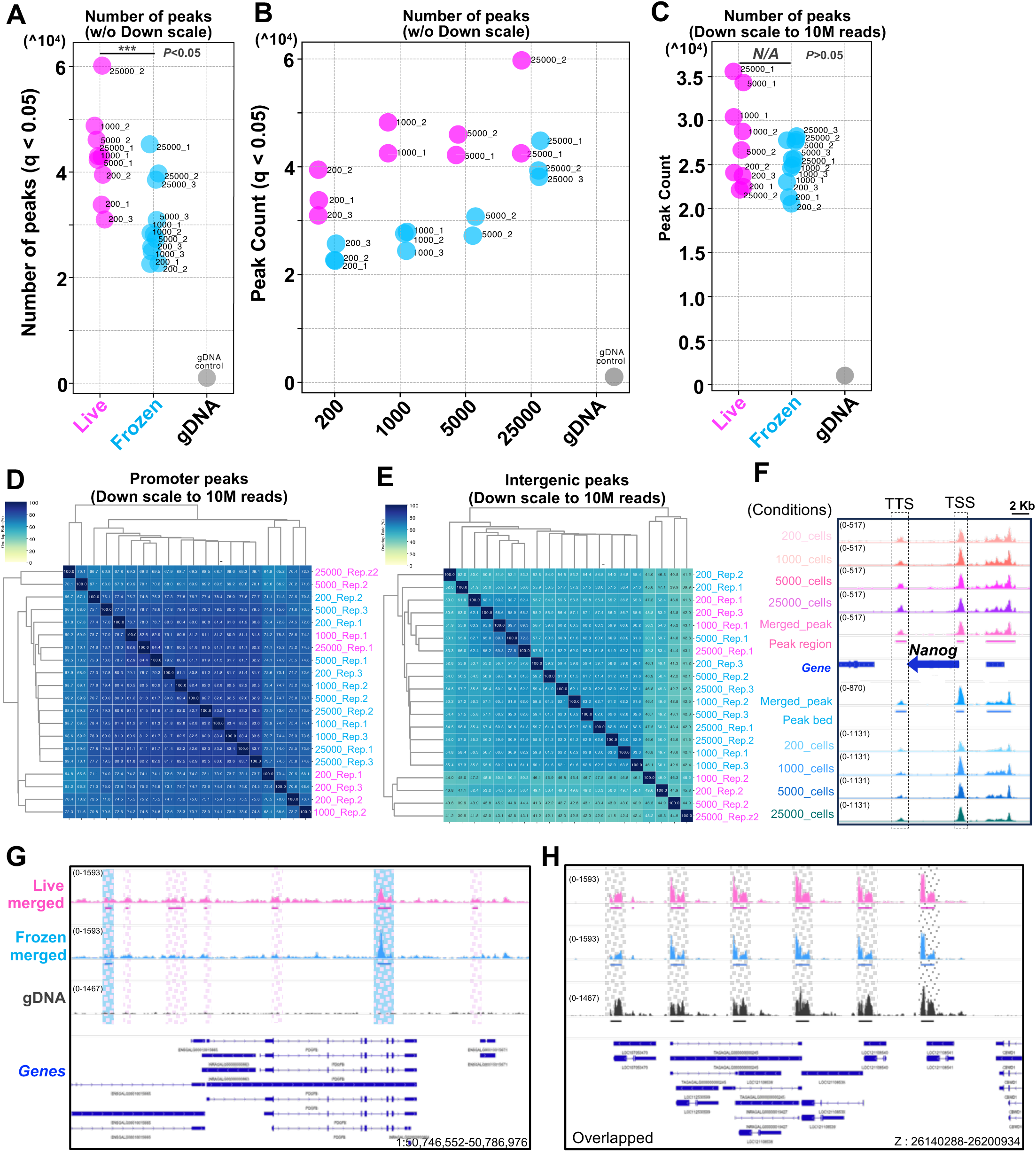
Omni-ATAC-seq from frozen PGCs reduced sensitivity for weak peaks in cryopreserved samples, whereas live and frozen samples still exhibit consistent promoter accessibility at strong peaks. (A) Number of peaks detected by MACS3 (--broad,--q 0.05) without read downscaling, across input sizes and preservation conditions. Peak numbers were significantly lower in frozen samples (blue) compared to live samples (magenta) (t-test, *P* < 0.05). (B) Relationship between input cell number and peak counts (without downscaling). Slight increases in peak counts were observed with higher input, regardless of preservation. (C) Number of peaks detected after normalizing all libraries to 10 million valid reads using scaling factors (no significant difference, *P* > 0.05). (D–E) Heatmaps showing pairwise peak overlaps after downscaling, for promoter-associated peaks (D) and intergenic peaks (E). Promoter peaks were highly concordant across all conditions, while intergenic peaks exhibited greater variability. (F) IGV tracks at *Nanog* locus show consistent peak detection across conditions and input levels at transcription start site (TSS) and transcription termination site (TTS), with merged replicates enhancing signal clarity. (G) Representative genome browser views showing accessible chromatin regions in live, frozen, and gDNA control samples. Live-specific low-intensity peaks are reduced or absent in frozen and gDNA samples. (H) Representative genome browser showed that frozen PGC ATAC-seq libraries fail to detect weak ACRs visible in live samples. Peaks in gDNA samples are enriched at interstitial telomeric sequences (ITSs), reflecting sequence preference of the Tn5 transposase.

We further assessed peak reproducibility by annotating all ATAC-seq peaks to genomic features (promoter, exon, intron, intergenic, and transcription termination site regions) and computing overlap rates. Promoter-associated peaks exhibited the highest reproducibility across all samples, with overlap rates of 70–85%, regardless of preservation method or cell number (Fig. 2D). Peaks located in intergenic, intronic, exonic, and TTS regions were more variable (∼65% overlap), likely due to the weaker and more heterogeneous nature of regulatory elements in these regions (Fig. 2E, Supplementary Fig. 1A–C).

To evaluate signal robustness at key regulatory loci, we visualized the detected ACRs at the *Nanog* promoter, a well-characterized marker of PGC identity and pluripotency (Choi *et al*., 2021). Consistent peaks were observed across all conditions, including cryopreserved 200-cell samples (Fig. 2F). To determine whether increased sequencing depth could restore detection of these weak peaks, we merged the reads in which after removing mtDNA and PCR-duplicated reads across the same conditions considered as the biological replicates (Fig. 2G). The merged datasets effectively recovered some of the weakly accessible regions, suggesting that deeper sequencing can mitigate the sensitivity loss associated with cryopreservation.

Lastly, we investigated potential sequence bias introduced by the Tn5 transposase using the gDNA control library, which can be utilized as a “blacklist.” Peak calling and motif enrichment analysis of this control yielded 1,023 peaks, disproportionately enriched in intergenic regions containing the telomeric (TTAGGG)n hexamer, likely corresponding to interstitial telomeric sequences (ITSs) (Alejandro D. Bolzán, 2017) (Fig. 2H, Supplementary Fig. 1D–J). The chicken genome is characterized by a distinctive karyotype comprising macro-and micro-chromosomes, and ITSs have been recognized as genomic signatures shaped by meiotic recombination and chromosomal rearrangements throughout its evolutionary history (I. Nadia et al., 2002). In light of such species-specific genomic features, assessing sequence bias using a gDNA negative control provides a valuable means to account for these intrinsic sequence properties when interpreting ATAC-seq results. This pattern most likely reflects inherent sequence preferences of Tn5 rather than genuine chromatin accessibility, underscoring the importance of distinguishing biological signals from technical artifacts in ATAC-seq datasets.

These results demonstrate that Omni-ATAC-seq applied to cryopreserved, low-input samples results in chromatin accessibility profiles largely comparable to those of live-cell samples, particularly in promoter regions. Nonetheless, higher sequencing depth or replicate integration may be required to capture low-abundance or weakly accessible regulatory elements in frozen material. Importantly, we identified technical artifacts occurring by intrinsic sequence preferences of the Tn5, which appear to be linked to species-specific chromosomal features such as ITSs in the chicken genome. By incorporating a gDNA-based negative control to detect and remove such artifacts, we provide a strategy to more accurately distinguish genuine biological signals from sequence bias. This approach is especially relevant when working with species that possess distinctive genomic architectures, and it expands the applicability of Omni-ATAC-seq to a wider range of rare or previously inaccessible cell populations in developmental and comparative genomics studies.

### Identification of PGC-Specific Accessible Chromatin Regions

We next ask whether it is possible to identify functional ACRs either PGC-specific or developmentally stage-specific, by integrative analysis with publicly available ATAC-seq datasets (Fig. 3A). To identify PGC-specific regulatory elements, instead of differential peak analysis we applied a binary comparison strategy, subtracting peaks shared with multiple somatic tissues to remove commonly accessible ACRs (Fig. 3A). We merged our datasets by input type (live or cryopreserved), each yielding 195–247 million mapped reads, so that the library volume became comparable to other tissue libraries (Supplementary Table 2). Peak calling produced 55,805 overlapping peaks between live and cryopreserved PGCs (Fig. 3B). Live-specific peaks (∼30,000) were far more numerous than cryopreserved-specific peaks (∼5,759), with shared peaks mainly in promoter regions and live-specific peaks enriched in intronic and intergenic regions, that is consistent with the previous results at figure 2 (Fig. 3C, Supplementary Table 3).

**Figure 3.**
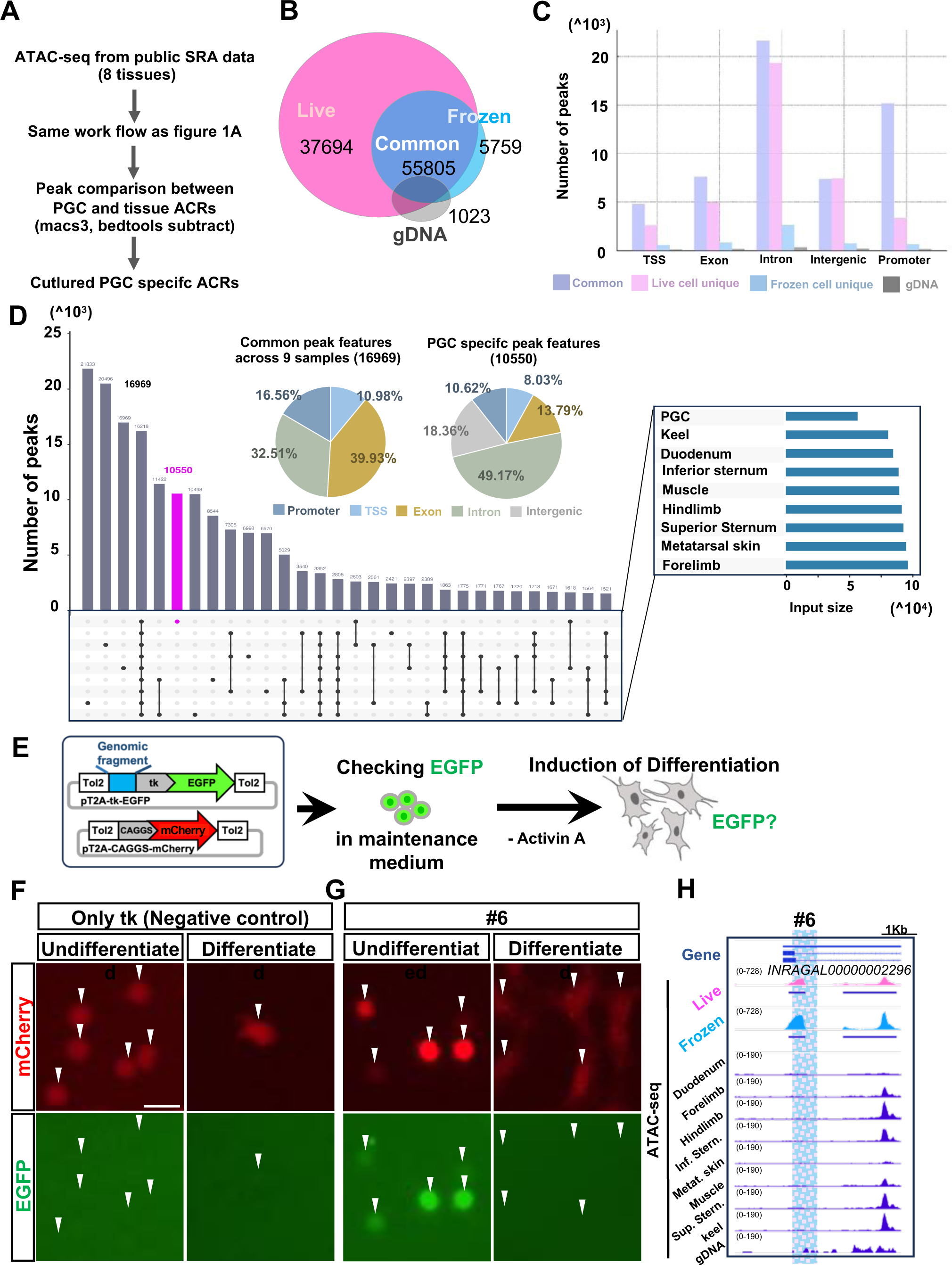
Comparison of PGC and somatic tissue ATAC-seq samples enabled the identification of cell-type-specific cis-regulatory elements, even from low-input frozen PGC ATAC-seq library. Schema of the analysis pipeline: ACRs identified in cultured PGCs were compared with published ATAC-seq datasets from somatic tissues to define PGC-specific ACRs. (B) Venn diagram showing the number of peaks detected in live and cryopreserved PGCs, and their overlap. A total of 55,805 peaks were shared between conditions, with live samples contributing a greater number of unique peaks. (C) Peak annotation of ACRs categorized by feature (Promoter, exon, intron, intergenic, TTS). Many populations of intronic and intergenic peaks were detected and biased in live samples. (D) Upset plot illustrating peak overlaps among 8 somatic tissue datasets. Bar plots indicate total peak numbers; pie charts show genomic feature distributions of commonly shared ACRs (left) and PGC-specific ACRs (right). (E) Schematic diagram of the transplantation assay with cultured PGCs. Cultured PGCs transfected with enhancer reporter constructs, CAGGS-mCherry, and CAGGS-T2TP plasmids were co-transfected in cultured PGCs. After the expression of EGFP was confirmed, the culture medium containing Activin was replaced with a differentiation medium (without Activin). (F) Negative control: tk-EGFP construct lacking an upstream enhancer (Only tk). (G) Enhancer construct #6. (H) IgV tracks showed genomic loci at ACR candidate #6. The peak tracks were shown ATAC-seq signals from merged live PGC (mazenta), merged frozen PGC (light blue) and individual 8 tissues (blue).

To define PGC-specific ACRs, we compared peaks shared by live and cryopreserved samples with a binary matrix from 30 public ATAC-seq datasets of 8 somatic tissues and embryonic organs (Fig. 3A). This comparison resulted in 10,550 PGC-specific ACRs (Fig. 3D, Supplementary Table 4), distributed across promoters (10.62%), exons (13.79%), introns (49.17%), intergenic (18.36%), and TTSs (8.03%). In contrast, 16,969 ACRs shared with somatic tissues were enriched in promoters (16.56%) and exons (39.93%), with no intergenic contribution. Overall, ACR numbers in PGCs were comparable to somatic tissues. These findings suggest PGC-specific expression is largely controlled by enhancers in non-coding regions rather than promoter-proximal sites, and intergenic ACRs at housekeeping genes are not uniformly accessible across tissues or cell types.

To identify functional PGC-specific enhancers, we selected 16 representative candidate regions from the >10,000 ACRs uniquely identified in PGCs by ATAC-seq (Supplementary Table 5). First, we filtered for peaks that were exclusively detected in PGCs and absent in somatic tissue samples. Next, we selected peaks that were consistently detected across various PGC input conditions (ranging from low to high input, including both live and frozen samples), aiming to evaluate whether functional enhancers could be robustly captured under suboptimal or practical experimental conditions. The selected candidates spanned a broad range of chromatin accessibility values (as determined by q-values and fold changes from peak calling) and were located within 2,500 bp of annotated gene bodies. Because ACRs can vary in size— from nucleosome-free regions (<100 bp) to regions spanning two or three nucleosomes—we excluded regions smaller than 50 bp and larger than 550 bp (approximately the width of >3 nucleosomes) to avoid excessively broad peaks that may reflect open chromatin regions without clear enhancer activity. This filtering strategy allowed us to include a representative range of accessibility profiles while minimizing potential artifacts. Ultimately, candidate enhancers were prioritized not only by accessibility rank but also by genomic context and technical feasibility, enabling a focused evaluation of their regulatory potential in PGCs.

### Functional Validation of Candidate PGC-Specific ACRs in Cultured Cells

To experimentally validate the enhancer activity of the selected PGC-specific ACRs, we performed a fluorescence-based reporter assay in cultured PGCs. Reporter constructs containing each candidate ACR upstream of a minimal tk promoter driving EGFP (Uchikawa *et al*., 2003) were co-transfected with a CAGGS-mCherry control plasmid to monitor transfection efficiency (Fig. 3E). Of the 16 candidate ACRs prioritized as described above, 14 genomic regions were successfully amplified by genomic PCR and cloned into the reporter vector. Two candidates could not be amplified and were excluded from further analysis (Supplementary Table 5).

The tk-EGFP vector lacking any upstream enhancer served as a negative control and produced only minimal EGFP signal (Fig. 3F). As a positive control, we used the Nanog promoter, a well-characterized PGC-expressed regulatory element, which drove robust EGFP expression (Supplementary Fig. 4), validating the sensitivity of our assay. Among the 14 tested constructs, the majority showed EGFP fluorescence above the basal level (#1, #3, #4, #5, #6, #8, #9, #12, #13, #14, and #15), and several candidates exhibited strong and consistent activity comparable to the Nanog promoter (Supplementary Fig. 2). The co-expression of mCherry in all samples confirmed successful transfection and enabled normalization of EGFP signal intensities, excluding variation in transfection efficiency as a confounding factor. The activities of the enhancer candidates were in strong agreement with the observation that the corresponding peak loci exhibited robust and representative signals across PGC datasets covering both low to high input amounts and live or frozen samples (Supplementary Fig. 3). These results provide direct experimental evidence that a substantial fraction of the ATAC-defined, PGC-specific ACRs, including those detectable even in frozen 200-cell datasets, function as *bona fide* enhancers in their native germ cell context.

### Loss of Enhancer Activity upon PGC Differentiation

To assess whether the enhancer activity of PGC-specific ACRs depends on the undifferentiated state, we examined EGFP expression following *in vitro* differentiation of PGCs. In the course of this study, we found that cultured PGCs spontaneously differentiate when transferred to a medium lacking Activin A (Fig. 3E). Under these conditions, PGCs that typically remain in suspension began to adhere to the culture dish and exhibited morphological changes characteristic of differentiation. Notably, the promoter activity of established germline genes such as *Nanog* and *DDX4* was markedly reduced in this condition, as evidenced by diminished EGFP and mCherry fluorescence in reporter assays (Supplementary Fig. 4). To our knowledge, this is the first report to define a culture condition that reliably induces spontaneous differentiation of chicken PGCs *in vitro*.

Using this differentiation system, we cultured PGCs transfected with enhancer reporter constructs in the absence of Activin A and monitored EGFP fluorescence before and after differentiation. Strikingly, EGFP signals driven by all enhancer candidates—including those with strong activity in undifferentiated PGCs such as #3, #4, #5, #6, #8, and #12—were abolished following differentiation (Fig. 3F-H; Supplementary Fig. 5). In contrast, mCherry fluorescence from the co-transfected control plasmid remained detectable in both undifferentiated and differentiated cells, confirming that the observed loss of EGFP was not due to differences in transfection efficiency or cell viability.

These findings indicate that the enhancer activity of the tested ACRs is tightly linked to the undifferentiated state of PGCs and becomes silenced upon differentiation. Together, these data support the interpretation that the tested ACRs represent germline-competent, state-dependent regulatory elements.

### PGC-Specific Enhancer Activity is Restricted to Early Development

Based on the identification of PGC-specific enhancers using ATAC-seq (Fig. 3), we next sought to refine the selection of candidate elements by incorporating stage-specific transcriptomic data from endogenous chicken PGCs (Fig. 4A). To determine whether stage-specific ACRs could also be detected, we integrated our ATAC-seq data with RNA-seq datasets from Ichikawa *et al*. <u>(Ichikawa et al. 2022)</u> differential gene expression analysis (DESeq2; baseMean > 10, q<0.01, |log₂FC|>2) between migrating PGCs at embryonic day 2.5 (E2.5) and gonad-resident PGCs at E6 identified 525 genes significantly upregulated at E2.5 and 270 genes upregulated at E6 (Fig. 4B, Supplementary Table 6). We then examined whether PGC-specific ACRs were located near these differentially expressed genes. Among the 21,892 filtered peaks, 434 were associated with genes highly expressed at E2.5, and 321 genes contained PGC-specific ACRs within 2,500 bp of their gene bodies (Fig. 4C). In contrast, loci of genes highly expressed at E6—such as *LDHA*—tended to show small, constitutive peaks shared across all tissues, lacking PGC-specific ACRs (Fig. 4C). This pattern likely reflects the fact that the cultured PGCs used for ATAC-seq were originally derived from E2.5 embryos.

**Figure 4.**
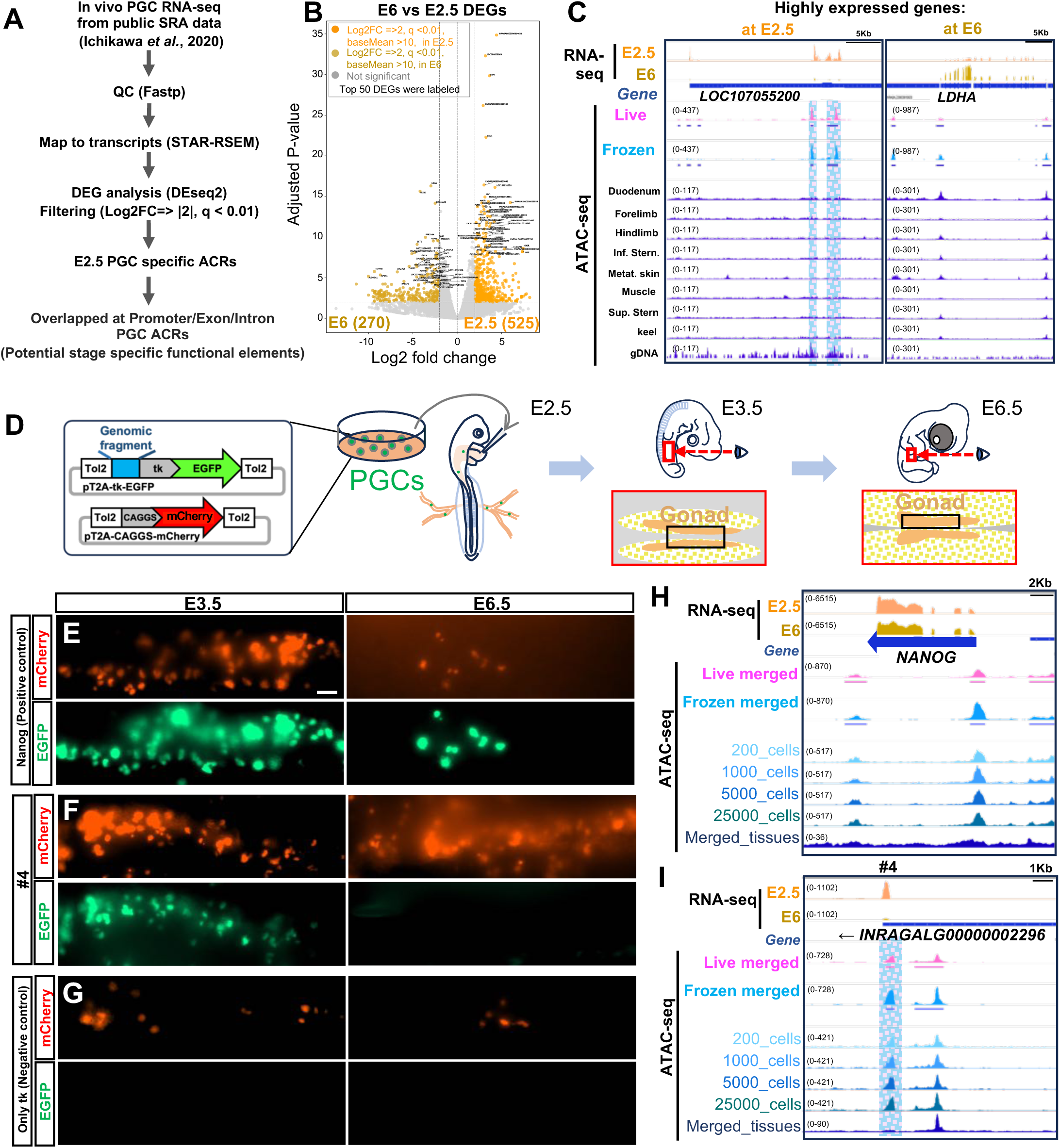
PGC-specific ATAC peaks exhibit enhancer activity *in vitro* and are downregulated upon differentiation. Schematic overview of the analysis workflow. (B) Volcano plots showing transcriptome re-analysis comparing *in vivo* PGCs at E2.5 and E6. Genes with absolute log₂ fold change > 2, q-value < 0.01, and baseMean > 10 were highlighted. Genes highly expressed at E2.5 are marked in light orange, and those at E6 in orange. The top 50 most significantly expressed genes are labeled. (C) IGV tracks showing genomic regions around ACR candidate #6. The top two tracks show transcript data from E2.5 and E6, respectively. ATAC-seq signals from merged live PGCs, merged frozen PGCs, and individual somatic tissues are displayed. The left panel shows high expression of LOC107055200 at E2.5, while the right panel shows high expression of LDHA at E6. (D) Schematic diagram of the transplantation assay. Cultured PGCs transfected with enhancer reporter constructs, CAGGS-mCherry, and CAGGS-T2TP plasmids were injected into the bloodstream of E2.5 chicken embryos. Embryos were harvested at either E3.5 or E6.5. (E-G) Fluorescence images of transplanted PGCs at E3.5 and E6.5, showing ventral views of the regions outlined in (D). (E) Positive control: Nanog promoter–EGFP construct. (F) Enhancer construct #4. (G) Negative control: tk-EGFP construct lacking an upstream enhancer (Only tk). Scale bars: 50 μm. All experiments were performed with n=5 embryos per condition. (H–I) IGV tracks showing transcripts from E2.5 and E6, along with ATAC-seq signals from merged live PGCs and frozen PGCs across input amounts. At the Nanog locus and INRAGALG00000002296, a highly expressed gene at E2.5, strong ATAC-seq signals were observed around the TSS and/or TTS. Nanog transcripts were detectable in both E2.5 and E6.

To assess stage-specific activity during PGC development, we performed *in vivo* transplantation of selected three candidate enhancers (#3, #4, #5) that had shown strong enhancer activity *in vitro* (Fig. 4D). Cultured PGCs transfected with the enhancer reporter constructs and a CAGGS-mCherry control plasmid were injected into the bloodstream of stage E2.5 chicken embryos. Transplanted embryos were collected and analyzed at two developmental time points: E3.5—when PGCs are migrating through the dorsal mesentery or have just arrived at the gonadal primordium—and E6.5, by which time PGCs have stably settled within the developing gonads. At E3.5, all tested enhancers exhibited detectable EGFP fluorescence, indicating that their activity is maintained during the migratory phase (Fig. 4 E– G; Supplementary Fig. 6). In contrast, at E6.5, EGFP signals were consistently absent, while mCherry fluorescence remained detectable, confirming successful transfection and survival of the transplanted PGCs. The peaks tested in these assays were not detected in any somatic tissue samples, yet showed highly robust accessibility in PGC ATAC-seq data. They were located near genes with high expression at E2 that markedly decreased by E6, providing consistent evidence for their PGC specificity and stage-dependent activity (Fig. 4H-I). Notably, a *Nanog* promoter construct continued to drive EGFP expression at both time points (Fig. 4E), supporting the conclusion that the loss of enhancer activity was not due to global transcriptional repression or cell death, but rather reflects stage-specific silencing of the tested PGC enhancers. These results demonstrate that the activity of these enhancers is not only cell type–specific, but also developmentally restricted—active during early migratory stages and silenced following PGC colonization of the gonadal niche.

In summary, we establish that Omni-ATAC-seq can generate high-quality chromatin accessibility profiles from as few as 200 cryopreserved chicken PGCs, enabling robust identification of cis-regulatory elements even under low-input conditions. This methodological advance removes a major technical barrier for epigenomic profiling of rare cell types and facilitates sample storage, transport, and cross-laboratory collaboration. Beyond its technical implications, our findings provide new opportunities to dissect the gene regulatory architecture that underlies germline specification and developmental stage–specific transcriptional programs in avian models. Given the unique experimental accessibility of chicken PGCs, this approach will not only accelerate germline biology research but also open the door to comparative and functional studies across diverse developmental systems.

## Materials and Methods

### Animals, staging, and animal care

Fertilized chicken (*Gallus gallus domesticus*, White leghorn) eggs were purchased from Yamagishi poultry farm (Mie, Japan) and from Nagoya University through the National Bio-Resource Project of the MEXT, Japan, respectively. Eggs were incubated at 38.5℃, and embryos were staged by Hamburger and Hamilton’s stage (Hamburger and Hamilton, 1951). All animal experiments were performed with the approval of the Institutional Animal Care and Use Committees at Kyushu University under the allowance number A25-189-1.

### Establishment of chicken cultured PGCs

Circulating PGCs, along with blood cells, were harvested from blood of HH15 chicken embryos and were cultured on non-coated (non-adhesive) plates in calcium-free DMEM (Gibco) diluted with water, supplemented with Fibroblast Growth Factor Chimera (Wako), <u>A</u>ctivin A (APRO Science group), and <u>c</u>hicken <u>s</u>erum (Biowest) (FAcs medium) according to previously described methods (Chen *et al*., 2019); (Whyte *et al*., 2015)). After one month of expansion, PGCs were cryopreserved at-80℃ in Bambanker (NIPPON Genetics) until further use in experiments.

### PGC frozen stock and ATAC-seq library preparation

ATAC-seq libraries from living and cryopreserved PGC were prepared based on the Omni-ATAC seq method with modifications as recently published ((Corces *et al*., 2017); (Kawaguchi *et al*., 2024)). Cultured PGCs were harvested, counted the cell density and aliquoted to the desired number of cells (200, 1000, 5000 and 25000) in individual DNA low binding tubes (Eppendolf, 0030108051). Cells were collected with centrifugation (350 G for 5 min at room temperature) (KUBOTA, 2410). For the cryopreservation, collected cells were individually resuspended into 50 μl of Banbanker (Nihon Genetics, CS-02-001). Tubes were slowly frozen using Mr.Frosty (Thermofisher, 5100-0001) and kept at-80°C until library preparation. When ATAC-preparations were conducted, cryopreserved cells were thrown at 37℃ for 5 min, and the cryopreserved samples were prepared together with living cultured PGC samples. Concurrently, living cultured PGCs were harvested as described above. In the cryopreserved and living cultured cells, 500 μl Lysis buffer 1 was added (10 mM Tris-HCl pH7.5, 3 mM MgCl2, 2 mM NaCl, x50 Protease inhibitor), and gently mixed and harvested (350 G for 5 min at 4°C) (KUBOTA, 2410). The cell pellets were resuspended in 50 μl Lysis buffer 2 (10 mM Tris-HCl pH7.5, 3 mM MgCl2, 2 mM NaCl, 0.1% Tween-20, 0.1% NP-40, 0.01% Digitonin, x50 Protease inhibitor) and incubated on ice for 15 min. 450 ul of Lysis buffer 3 (10 mM Tris-HCl pH7.5, 3 mM MgCl2, 2 mM NaCl, 0.1% Tween-20) was added to the cells and mixed gently. Immediately, cells were spun down (350 G for 5 min at 4°C) and pellets were resuspended with 25 μl of Tn5 Transposase reaction for the input of 200 and 1000 cells, and 50 μl Tn5 Transposase reaction for the input on 5000 and 25000 cells (Transposase reaction: 12.5 μl of the 2x Transposase buffer (20 mM Tris-HCl (pH7.5), 10 mM MgCl2, 20% DMF), 16.5 μl of PBS, 0.5 μl of 10% Tween-20 (0.1% f. c.), 0.5 μl of 1% Digitonin (0.01% f. c.) and 2.5 μl of assembled in-house Tn5 (0.5 μg of in-house Tn5 stock was diluted in 1:10)). Tagmentation proceeded at 37°C in water-bath for 1 hour with occasional agitation. The reaction was stopped by adding a 5 times volume of PB (QIAGEN, 19065) and vertexing for 30 seconds. Tn5 treated DNA was purified with MinElute PCR purification Kit (QIAGEN, 28004) and eluted with 20 μl of EB buffer. 10 μl of purified DNA solution was used for library amplification, and final amplification cycles were defined with intermediate qPCR as described in the original ATAC-seq method (Buenrostro et al., 2013). Nonetheless, all libraries were amplified with 9-16 total PCR cycles in the deserved number. Final libraries were amplified with NEBNext® High-Fidelity 2XPCR Master Mix (NEB, M0541)) with Nextera Unique Dual index (UDI) adapter plate A (illumina, #1000000002694). After PCR amplification, the libraries were purified with magnetic beads for DNA isolation. Removing Adapters were prepared following the manufacturing protocol. Sequencing was performed using Nova-seq6000 PE150 at the third party.

### Computational analysis and data visualization Reference genome and annotation versions

Throughout the analysis, the reference genome assembly bGalGal1.mat.broiler.GRCg7b (NCBI accession; GCF_016699485.2) and gene annotation from Gallus Enriched Gene Annotation were used as reference gene model (Degalez *et al*., 2024) (https://gega.sigenae.org).

### ATAC-seq processing and peak analysis

Trimming of Nextera adapters and filtering of low-quality reads were performed using Fastp (v0.23.2) with the parameters (-q 15-n 10-t 1-T 1-l 20) (Chen *et al*., 2018). Bowtie2 (v2.5.4) (Langmead *et al*., 2019) was used for read mapping with the parameters (--no-mixed--no-discordant-X 1000). SAMtools (v1.17) (Danecek *et al*., 2021) was used to convert SAM files to BAM files, and sorting and indexing were performed with default settings. Picard (v3.0.0) (https://broadinstitute.github.io/picard/), was used for duplicate removal and fragment size distribution analysis with the MarkDuplicates and CollectInsertSizeMetrics tools. Duplicate-removed BAM files were further used to assess mtDNA and unmapped reads using SAMtools with the parameters (view-f 4). Peaks were detected with MACS3 peakcall (v3.0.0a6), and used the--broad option to account for the nature of accessible regions spanning multiple nucleosomes with the parameters (-f BAMPE-q 0.05-g 1053332251--broad) (Zhang *et al*., 2008). Peaks were ranked based on p-values and q-values obtained from MACS3. Peak comparisons were performed using Bedtools (v2.31.0) (Quinlan and Hall, 2010), and annotations were carried out using annotatePeaks.pl from HOMER (v4.9.1) <u>(Heinz et al. 2010</u>; http://homer.ucsd.edu/homer/). To identify potential PGC-specific ACRs, we focused on ACRs detected in both live and cryopreserved samples. For binary comparisons, peaks across all samples were intersected using Bedtools (v2.31.0). To exclude gDNA-derived peaks from the PGC ATAC-seq data, overlapping peaks were identified with Bedtools intersect with the parameters (-u-f 0.5), and subsequently subtracted. For the data visualization, generating bigwig files, correlation heatmaps were carried out with multiBamSummary, plotCorrelation and bamCoverage by DeepTools (v3.1.2) with the parameters (--binSize 10 –normalizeUsing RPKM--smoothLength 30--effectiveGenomeSize 1053332251). The upsert plot was generated by intervene (v0.6.5) (Quinlan and Hall, 2010; Khan and Mathelier, 2017). For motif analysis, 1,023 peaks from gDNA tagmentation and 1,023 ACRs from merged live PGC samples (randomly selected using Bedtools) were converted to FASTA format and analyzed for enriched motifs using MEME (Bailey *et al*., 2015).

### RNA-seq analysis

RNA-seq datasets were obtained from publicly available data from K. Ichikawa et al. (2022), consisting of four biological replicates each of *in vivo* PGCs at E2.5 and E6. The datasets were obtained from the Gene Expression Omnibus under accession number GSE188689. Trimming of Nextera adapters and filtering of low-quality reads were performed using Fastp (v0.23.2) with the parameters (-q 15-n 10-t 1-T 1-l 20-w 16) (Chen *et al*., 2018). Mapping and generation of count matrices were conducted using the STAR-RSEM pipeline with default parameter (Li and Dewey, 2011; Dobin *et al*., 2013). Differential expression analysis was carried out using PyDESeq2 (cutoff: baseMean >10, *q*>0.01) (Muzellec *et al*., 2023). For visualization, fastq files processed by Fastp were mapped using STAR to generate “Aligned.toTranscriptome.bam” files. Final bigWig files were generated using deepTools (v3.1.2) with the parameters (--binSize 10--normalizeUsing RPKM--smoothLength 30--effectiveGenomeSize 1053332251).

### Tn5 transposase purification and assembly

The pTXB1-Tn5 expression plasmid (Addgene, #60240) was digested with XbaI and BamHI, and fragment of Tn5-intein-CBD was subcloned into the pE21a (+) vector (NovoPro, V011023) as described by Sato et al., (Sato *et al*., 2019). Tn5 was expressed and purified as described (Picelli *et al*., 2014; Sato *et al*., 2019), with the following modification. The *Escherichia coli* Rosetta 2 cells (DE3) carrying the Tn5 expression plasmid were grown in 3 linters (1.5 L x 2) of Luria-Broth (LB) medium containing 100 µg/mL ampicillin and 34 µg/mL chloramphenicol at 37 °C until the optical density reached 0.9 at 600 nm. The cell cultures were cooled to 10 °C and further cultivated for 15 h after addition of isopropyl β-D-1-thiogalactopyranoside (IPTG) to a final concentration of 0.25 mM to induce protein expression. Following a further 4-hour cultivation at 23 °C, the cells were harvested by centrifugation and resuspended in 50 mL of Lysis buffer (20 mM HEPES-KOH pH 7.2, 0.8 M NaCl, 1 mM EDTA, 0.2% (w/v) Triton X-100, 10% (w/v) glycerol) supplemented with cOmplete protease inhibitor cocktail (EDTA-free, Merck). The cells were disrupted by sonication and clarified by centrifugation at 28,000 g at 4 °C for 45 min. 50 mL of the cell lysate (supernatant fraction) was gradually mixed with 5 mL of 10% (v/v) polyethyleneimine (pH 7.5) after which the mixture was incubated at 4 °C for 20 minutes. It was then clarified by centrifugation at 20,000 g at 4 °C for 20 min. The resulting supernatant fraction was mixed with 85 mL of Lysis buffer containing protease inhibitor and then applied to a 17 mL bed volume of chitin resin (New England BioLabs). The resin was then washed with 250 mL of Lysis buffer followed by 20 mL of lysis buffer containing protease inhibitor and 100 mM dithiothreitol (DTT). The resin was resuspended in the same buffer and incubated at 4 °C for 36 hours. The eluate was dialyzed against D1 buffer (25 mM HEPES-KOH pH 7.2, 0.2 M NaCl, 0.2 mM EDTA, 0.2% (w/v) Triton X-100, 20% (w/v) glycerol) for 15 hours. The dialyzed fraction was applied to a 1 mL Resource S column (Cytiva). The column was developed with a linear gradient of 0.2 to 1 M NaCl in H buffer (20 mM HEPES-KOH pH 7.2, 0.2 mM EDTA, 20% (w/v) glycerol) totaling 15 mL. The peak fractions were collected and dialyzed against D2 buffer (100 mM HEPES-KOH pH 7.2, 0.2 M NaCl, 0.2 mM EDTA, 10% (w/v) glycerol) at 4 °C for 15 hours and then mixed with glycerol equivalent to 0.8 volume of the dialyzed fraction. The aliquots were snap-frozen in liquid nitrogen and stored at-80 °C. Oligo and Tn5 assembly was produced according to the publication (Picelli *et al*., 2014). Tn5 assembled with Oligo was kept at-20C up to one month.

## Plasmid constructions

### pT2A-tk-EGFP

The pT2AL200R150G vector (Kawakami, 2005) was digested with XhoI and BglII. A DNA fragment containing multiple cloning sites (MCS), the tk promoter, EGFP coding sequence, and polyadenylation (polyA) sequence was amplified by PCR from ptkEGFP (Uchikawa *et al*., 2003; Kawakami, 2005). This fragment was then recombined with the digested the pT2AL200R150G vector using the In-Fusion^®^ HD cloning kit (TAKARA).

### pT2A-CAGGS-mCheery

The pT2A-CAGGS-MCS1-2A-MCS2 vector (Saito *et al*., 2022) was digested with NotI and MluI. A mCheery coding sequence was amplified by PCR. This fragment was ligated into the digested the pT2A-CAGGS-MCS1-2A-MCS2 by Mighty Mix DNA Ligation Kit (TAKARA).

### pT2A-Genomic fragment:tkEGFP

The genomic region of interest was isolated and amplified by PCR from the genome of BL-E strain chickens maintained at Nagoya University. The amplified fragments were digested with XhoI-EcoRV, XhoI, or EcoRV-BglII and subsequently ligated into XhoI-EcoRV or EcoRV-BglII digested pT2A-tkEGFP.

### pT2A-Nanog:EGFP

The pT2AL200R150G vector was digested with XhoI and BglII. A DNA fragment containing multiple cloning sites (MCS), EGFP coding sequence, and polyadenylation (polyA) sequence was recombined with the digested the pT2AL200R150G vector using the In-Fusion^®^ HD cloning kit (TAKARA), resulting in the pT2A-MCS-EGFP plasmid. A 4,000 bp upstream region from the Nanog start codon was isolated and amplified by genomic PCR from BL-E strain chickens. This fragment was ligated into the MluI-EcoRV site of pT2A-MCS-EGFP.

### pT2A-MVH(DDX4):mCheery

The pT2AL200R150G vector was digested with XhoI and BglII. A DNA fragment containing multiple cloning sites (MCS), mCheery coding sequence, and polyadenylation (polyA) sequence was recombined with the digested the pT2AL200R150G vector using the In-Fusion^®^ HD cloning kit (TAKARA), resulting in the pT2A-MCS-mCheery plasmid. An approximately 2,000 bp upstream genomic region from the start codon of the mouse VASA homolog (MVH), kindly provided from Dr. Bertrand Pain, was amplified and recombined with the MluI-EcoRV-digested pT2A-MCS-mCheery vector using the In-Fusion^®^ HD cloning kit. KOD One^®^ (Toyobo) was used for all PCR reactions. Primers and amplified genomic fragments are listed in Supplementary Table 5.

### Plasmid transfection

A total of 5 x 10^4^ cultured PGCs were washed with OPTI-MEM (Gibco) and placed into a 96-well coated plate containing 100 μl of FAot medium (adding Ovo-transferrin instead of chicken serum) (Whyte *et al*., 2015) without heparin and antibiotics (penicillin, streptomycin, and amphotericin) for 3 hrs. A transfection mixture containing 0.2 μg of total plasmids and 0.5 μl of Lipofectamine 2000 (ThermoFisher Scientific) in 50 μl of OPTI-MEM was then added to the wells. The ratio of plasmid concentrations used was 3:1 for reporter vector: CAGGS-mCherry vector, and an appropriate amount of Tol2 transposase vector (Sato *et al*., 2007) was co-transfected accordingly. 6 hrs after transfection, the medium was replaced with conventional FAcs medium (maintenance medium) containing heparin and antibiotics.

### Induction of PGC differentiation

After transfection, PGCs were cultured for one week in a maintenance medium. Subsequently, the cells were transferred to tissue-culture treated plates containing differentiation medium, which was prepared by omitting Activin A from the standard maintenance medium. The medium was refreshed by replacing half of the volume with fresh differentiation medium every two days.

### PGC transplantation

Transfected PGCs were resuspended in Opti-MEM (Gibco) at a concentration of 5,000 cells/µl and injected into host chicken embryos at HH15-16. The embryos were incubated at 38.5℃ and collected after 1 or 4 days of development, corresponding to E3.5 and E6.5, respectively.

### Imaging

Fluorescence images of cultured cells were acquired using an inverted microscope (IX83, Olympus) equipped with an sCMOS camera (Zyla-4.2 plus, ANDOR) and operated with cellSens software (Olympus). Embryo images were captured using a cooled CCD camera (ORCA-R2, HAMAMATSU Photonics) mounted on a macro zoom microscope (MVX10, Olympus) and controlled by High Speed Recording (HSR) software (HAMAMATSU Photonics).

## Author contributions

AK and DS: Study design, data analysis, investigation, methodology, writing-original draft, writing-final manuscript and funding acquisition. MI: Data analysis, *in vitro* culture experiments, investigation. HI, YM, MS, YN, SK and KI: Data analysis, methodology and discussion. All authors reviewed and approved the manuscript.

## Acknowledgment

We thank Dr. Haruki Ochi from Kobe university for the critical reading of the manuscript and comments. Dr. Yohey Ogawa from National Institute of Genetics, and Dr. Yuya Okusaki and Dr. Kenichi Nishijima from University of Nagoya for the fruitful discussion. This work is supported by the following foundations, AK, DS and KI: “Strategic Research Projects” grant from ROIS (Research Organization of Information and Systems). AK: Grant-in-Aid for Scientific Research KAKENHI (C) 23K05798, Grant-in-Aid for Transformative Research Areas (A) 25H02584 and The Naito Science & Engineering Foundation. DS: Grant-in-Aid for Scientific Research KAKENHI (B) 23K23897.

## Data and resource availability

All new sequence reads were deposited in the DNA Data Bank of Japan (DDBJ) Sequence Read Archive under BioProjectID under accession PRJDB37400, in the Genome Sequence Archive under Accession SAMD01620406-SAMD01620426. All data generated in this study are available within the article and its Supplementary Data files.

Supplementary Table 1: Details of ATAC-seq libraries prepared in this study

Supplementary Table 2: SRA data details and accession numbers utilized in the analysis

Supplementary Table 3: Annotated peak features using HOMER annotatePeaks.pl

Supplementary Table 4: Genomic location of PGC-specific ACRs

Supplementary Table 5: ACR information validated in vivo and in vitro

Supplementary Table 6: Differential expression genes in E2.5 vs E6 In vivo PGCs

---

**Supplementary Figure 1:**
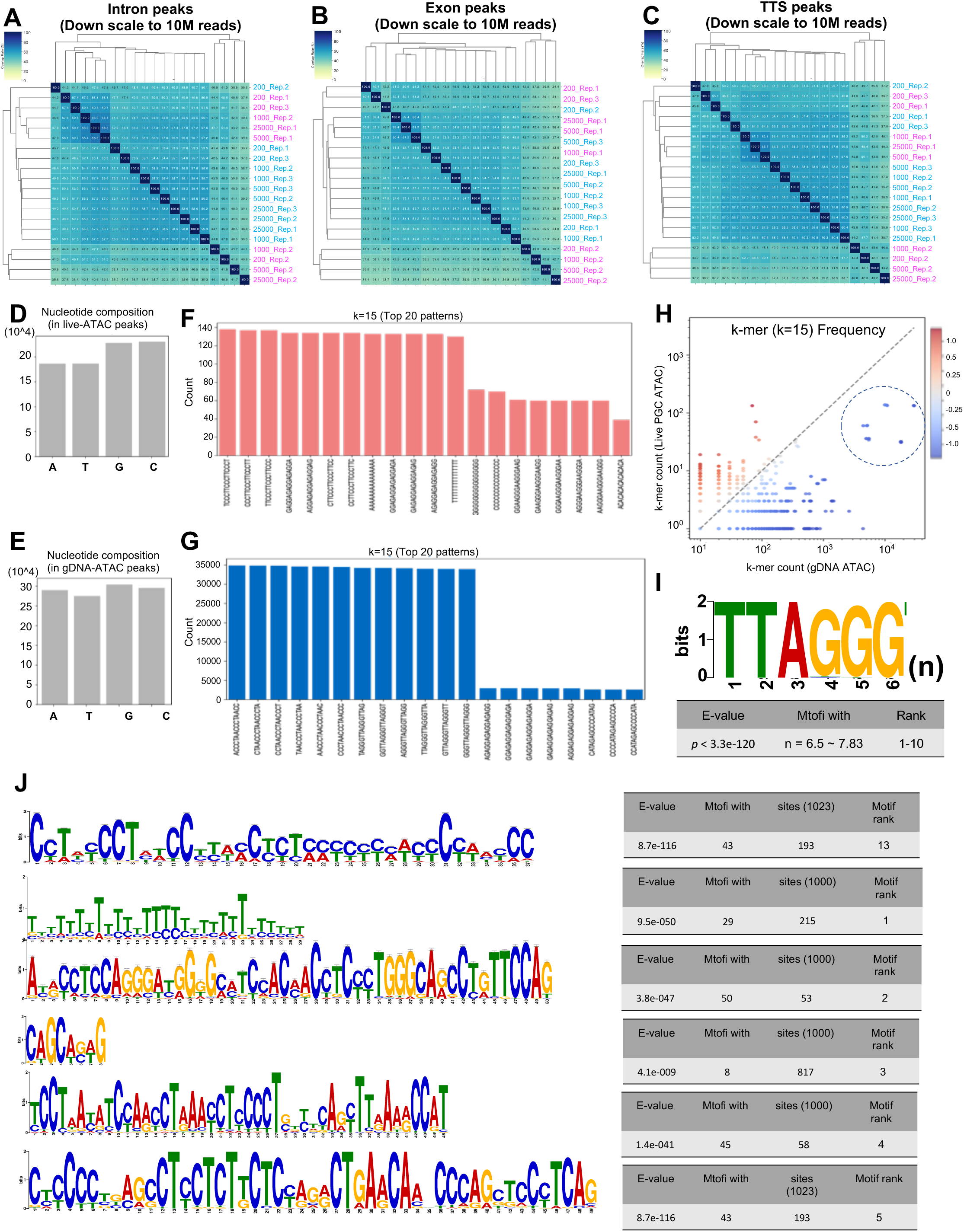
Evaluation of peak overlap, sequence bias, and motif enrichment in control and experimental samples. (A–C) Heatmaps showing the percentage of peak overlap across samples for intronic and exonic regions, and transcription termination sites (TTS) based on HOMER annotations. These features exhibited lower reproducibility (∼60–65%) compared to promoter peaks, regardless of input condition. (D-E) Nucleotide composition analysis of live-cell peaks and gDNA control peaks. For the comparison, 1,023 peaks were randomly sampled from the live-cell dataset to match the number of peaks identified in the gDNA library. The live-cell peaks showed a mild GC bias, likely reflecting promoter enrichment. In contrast, gDNA peaks displayed no strong nucleotide bias. (F-G) K-mer analysis (k=15) revealed strong sequence preference in the gDNA peaks, indicative of Tn5 transposase sequence bias, which was not observed in live-cell peaks. (H) Frequency distribution of control-biased sequences found in gDNA peaks, confirming that gDNA libraries tend to capture non-specific, sequence-driven Tn5 activity. (I) MEME motif analysis of gDNA-derived peaks revealed highly enriched tandem repeats of the canonical telomeric sequence (TTAGGG)n, with 6.5 – 7.8 repeats per motif. These sequences were frequently located in intergenic regions and likely correspond to interstitial telomeric sequences (ITSs), emphasizing that peaks from gDNA reflect intrinsic sequence preferences rather than chromatin accessibility. (J) In contrast, MEME analysis of live-cell ATAC-seq peaks showed no significant enrichment for repetitive telomeric motifs, further supporting that accessible chromatin profiles in live samples reflect true biological regulation.

**Supplementary Figure 2:**
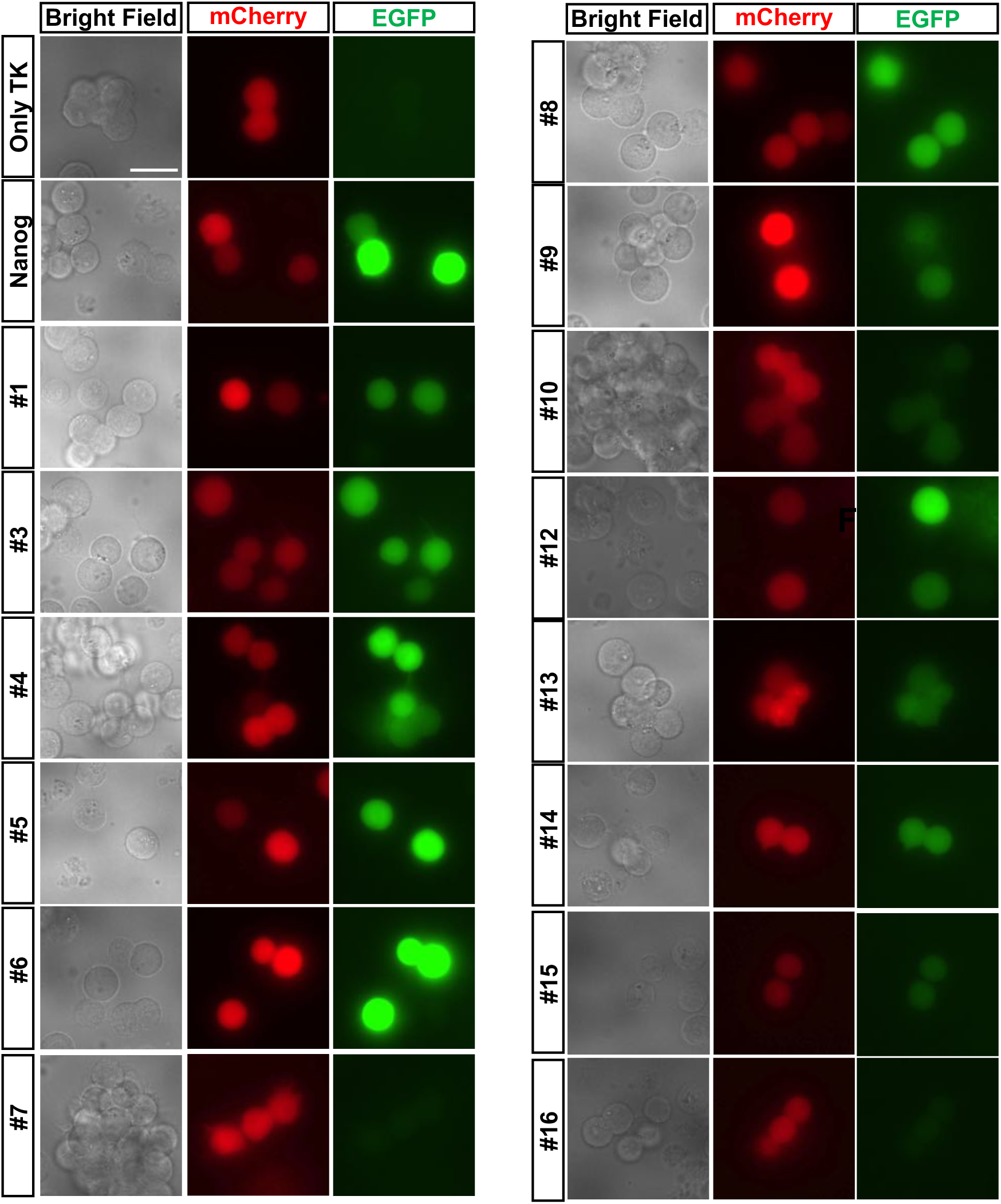
Fluorescence images of PGCs transfected with individual enhancer reporter constructs. Cultured PGCs were co-transfected with individual Tol2-based enhancer– EGFP reporter constructs, CAGGS-mCherry, and CAGGS-T2TP plasmids. Each genomic fragment was cloned upstream of a minimal tk promoter driving EGFP. Images were taken under maintenance conditions (+Activin A). For each construct, EGFP and mCherry fluorescence are shown alongside bright-field images. The tk-EGFP vector lacking an enhancer (Only tk) served as a negative control. The Nanog promoter–EGFP construct was included as a positive control. Scale bars: 20 μm. All experiments were performed with n=5 embryos per condition.

**Supplementary Figure 3:**
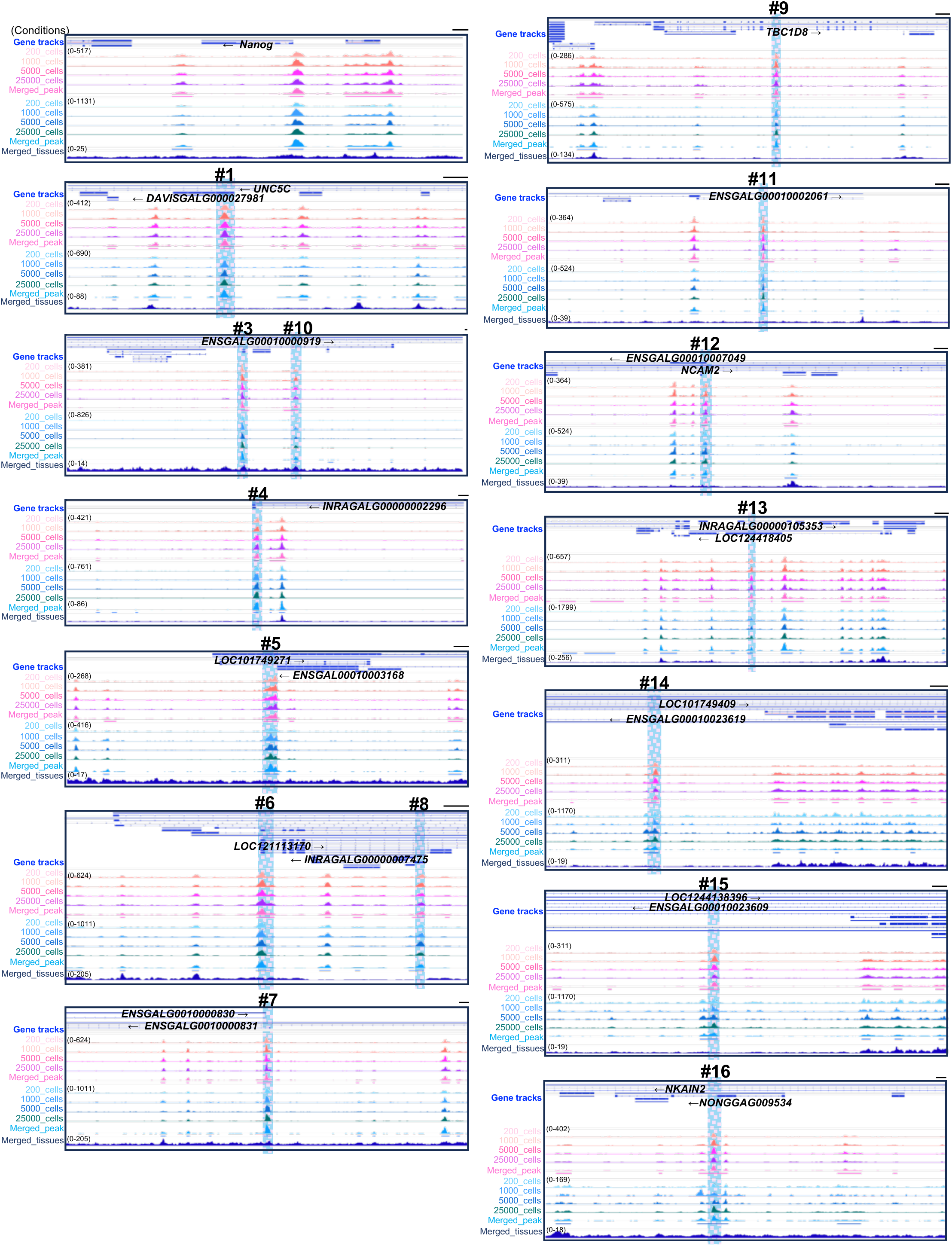
Genome tracks of PGC ACRs which were validated in this study. Representative IgV tracks at the genomic loci of 14 ACR were identified as uniquely accessible in PGCs, and its absence from somatic tissue datasets. Tracks showed ATAC-seq signal intensities for live and cryopreserved PGCs (low to high input amounts), genomic DNA controls, and merged somatic tissue ATAC-seq libraries (color-coded by sample type). Gene models (bottom) indicate annotated loci near each candidate region. Each candidate region exhibits a robust accessibility peak in PGCs, with consistent signal across live and frozen conditions, and no detectable signal in somatic tissues. Scale bar: 1 Kb.

**Supplementary Figure 4:**
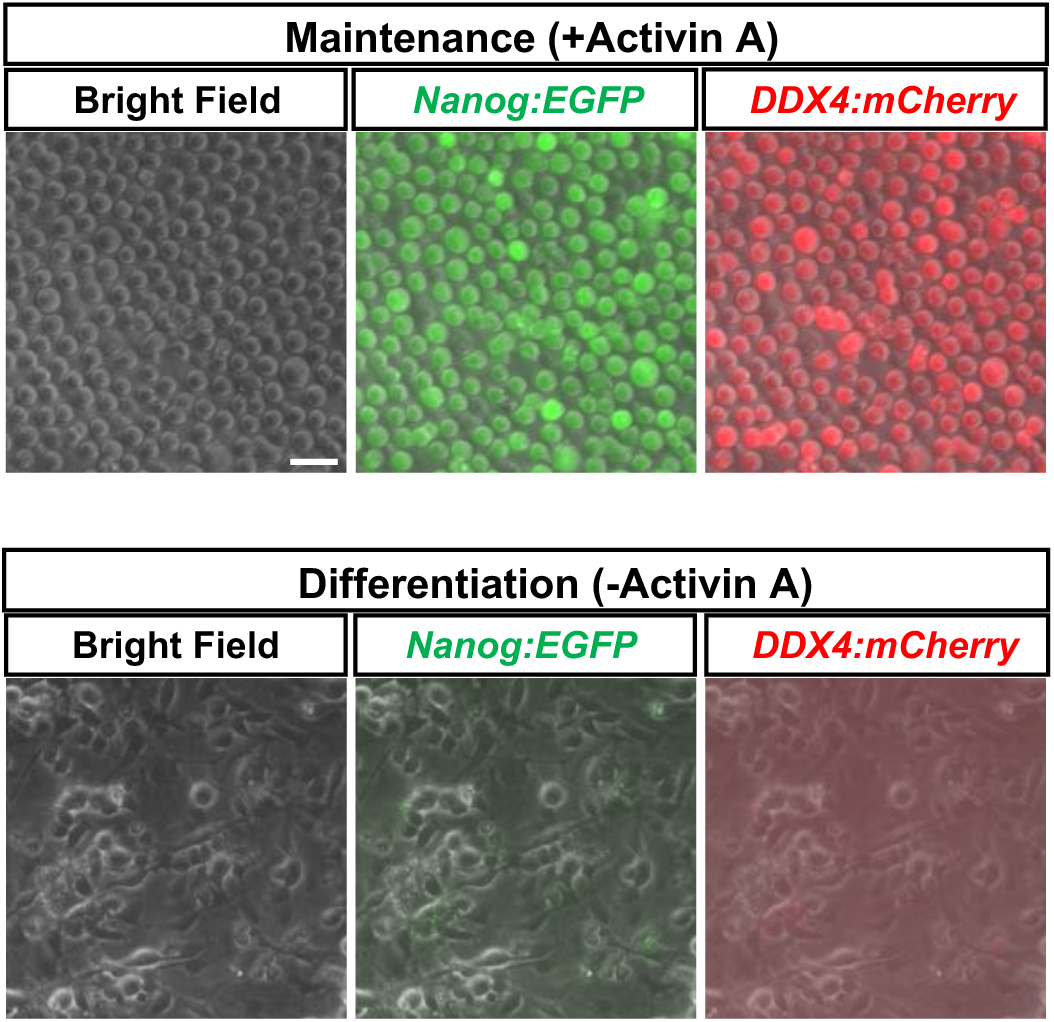
Loss of Nanog-and DDX4/MVH-promoter reporter signals upon differentiation of cultured chicken PGCs. Cultured chicken PGCs were co-transfected with a Nanog promoter–EGFP reporter (Nanog:EGFP) and a DDX4 promoter–mCherry reporter (DDX4:mCherry), then maintained in maintenance medium (+Activin A) or switched to differentiation medium (−Activin A). Under maintenance conditions, both Nanog:EGFP and DDX4:mCherry signals were readily detected, whereas under differentiation conditions both signals were abolished. Bright-field images are shown for reference. Scale bar: 30 µm.

**Supplementary Figure 5:**
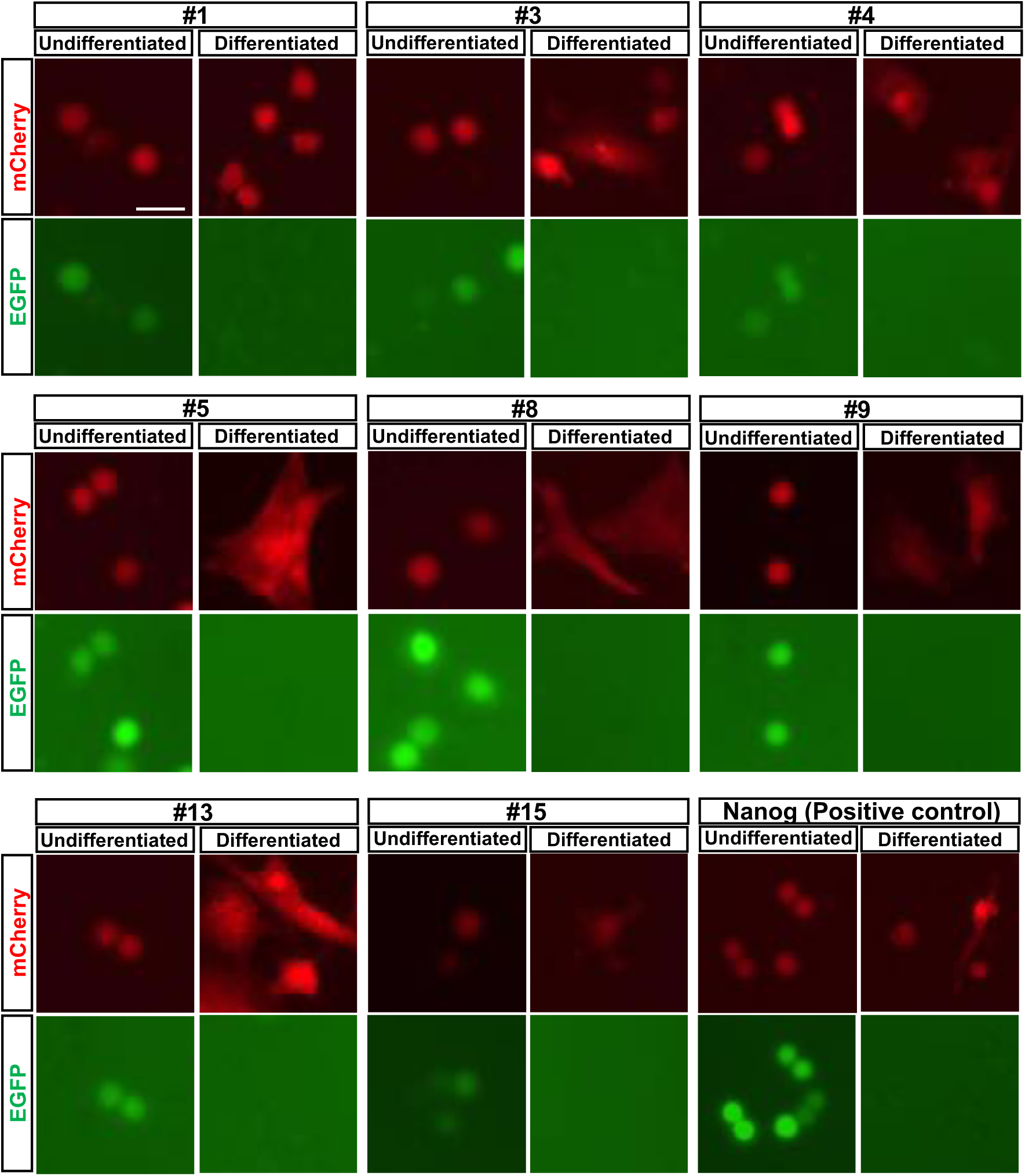
Fluorescence images of enhancer-reporter PGCs before and after *in vitro* differentiation. PGCs were transfected with individual Tol2-based enhancer–EGFP reporter constructs, along with CAGGS-mCherry and CAGGS-T2TP plasmids, and cultured under maintenance conditions (+Activin A) or differentiation conditions (−Activin A) for 3 weeks. For each construct, EGFP and mCherry fluorescence were observed and compared between conditions. All enhancer constructs that exhibited EGFP fluorescence under maintenance conditions showed reduced or undetectable EGFP signal upon differentiation. mCherry signal remained detectable across both conditions. Scale bars: 25 μm. All experiments were performed with n=5 embryos per condition.

**Supplementary Figure 6:**
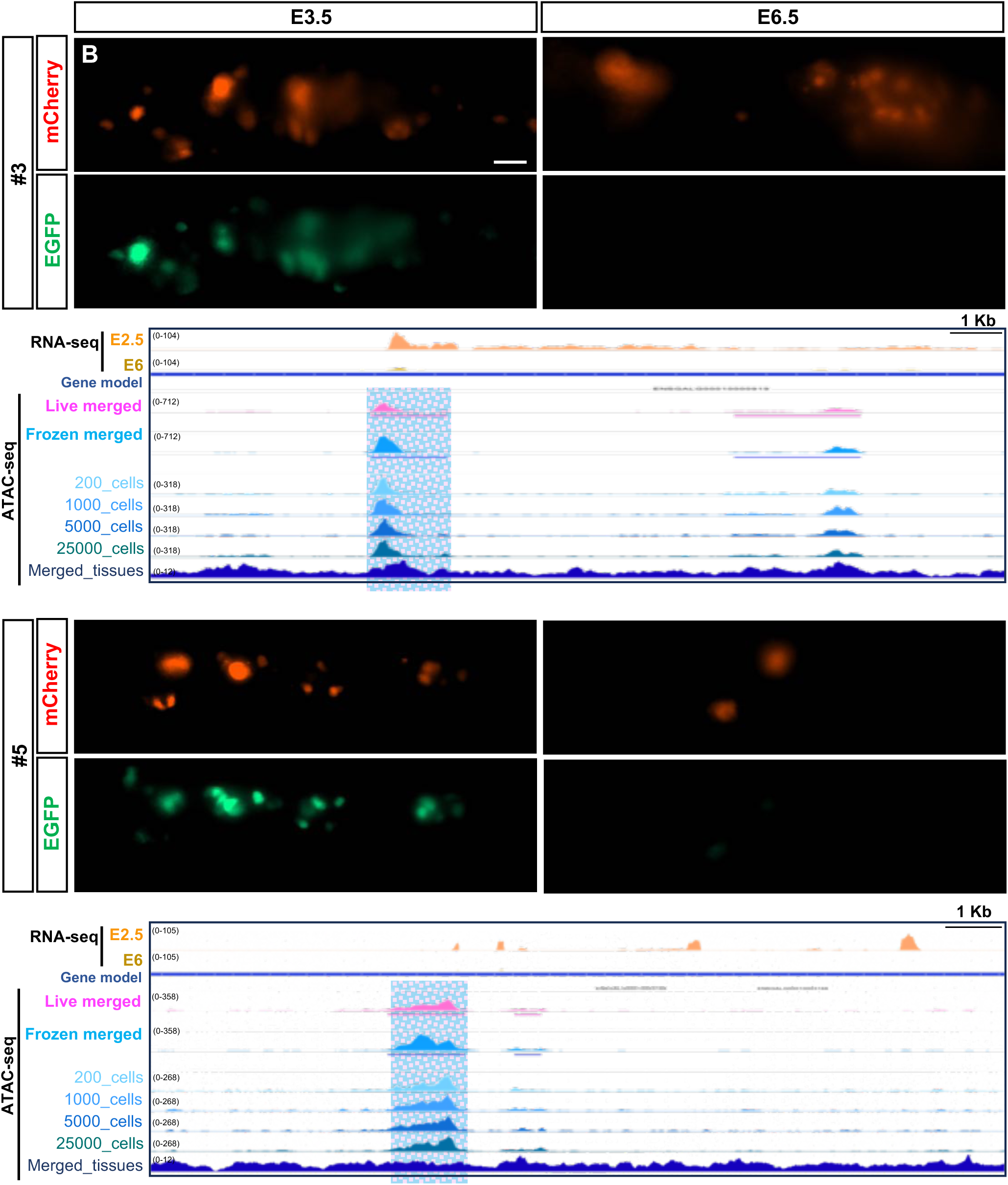
Fluorescence images of transplanted PGCs expressing individual enhancer constructs at E3.5 and E6.5. Cultured PGCs were transfected with individual Tol2-based enhancer–EGFP reporter constructs, CAGGS-mCherry, and CAGGS-T2TP plasmids, and transplanted into the bloodstream of E2.5 chicken embryos. Fluorescence images were acquired at E3.5 and E6.5, showing ventral views of the regions where PGCs were located. For each enhancer construct, EGFP and mCherry fluorescence are shown. Scale bars: 50 μm. All experiments were performed with n=5 embryos per condition. Individual IGV tracks displaying the corresponding genomic loci and ATAC-seq peaks from merged live PGCs and frozen PGCs across a range of input amounts (low to high).

